# Manifold Transform by Recurrent Cortical Circuit Enhances Robust Encoding of Familiar Stimuli

**DOI:** 10.1101/2025.03.02.641067

**Authors:** Weifan Wang, Xueyan Niu, Liyuan Liang, Tai-Sing Lee

## Abstract

A ubiquitous phenomenon observed along the ventral stream of the primate hierarchical visual system is the suppression of neural responses to familiar stimuli at the population level. The observation of the suppression of the neural response in the early visual cortex (V1 and V2) to familiar stimuli of size that are multiple times larger in size than the receptive fields of individual neurons reflects the plausible development of recurrent circuits for encoding these global stimuli. In this work, we investigated the neural mechanisms of familiarity suppression and showed that an excitatory recurrent neural circuit, consisting of neurons with small and local receptive fields, can develop to encode specific global familiar stimuli robustly as a result of familiarity training. This Hebbian learning based model attributes the observed familiarity suppression effect to the sparsification of the population neural code for the familiar stimuli due to the formation of image-specific local excitatory circuits and competitive normalization among neurons, leading to the paradoxical neural response suppression to the familiar stimuli at the population level. We explored the computational implications of the proposed circuit by relating it to the sparse manifold transform. The recurrent circuit, by linking spatially co-occurring visual features together, compresses the dimensions of irrelevant variations of a familiar image in the neural response manifold relative to the dimensions for discriminating different familiar stimuli. The computation can be considered as a globally non-linear but locally linear manifold transform that orthogonalizes the slow modes of network dynamics relative to the subspace of irrelevant stimulus variations, resulting in increased robustness of the global stimulus representation against noises and other irrelevant perturbations. These results provide testable predictions for neurophysiological experiments.

**Author summary:** In this research, we explored how the brain can become more efficient at processing familiar visual information. When we repeatedly see something, our brain’s response to it changes. In response to familiar stimuli, neurons across the different visual areas of the mammalian visual system become more selective and their overall activities decrease. We developed a computational model to investigate why this happens and what functional advantages these mechanisms might provide. We discovered that familiarity leads to the development of a more efficient and robust neural representation of what we see. It allows us to rapidly and robustly recognize our friend’s face despite changes in lighting conditions, view angle, or facial expression. Our model showed that through repeated exposure, the brain’s neural circuits, even in the early stages of visual processing, rapidly adapt and organize themselves to focus on important and consistent features in our visual environment while becoming less sensitive to irrelevant variations, and distractions.

## 1 Introduction

Familiarity suppression refers to a phenomenon observed in the inferotemporal cortex (ITC) [1, 2, 3, 4, 5, 6, 7] and more recently in early visual cortex [8] that repeated exposure to a set of familiar visual stimuli will lead to the suppression of neural responses to these stimuli, particularly in the later part of the temporal responses. There is evidence in the inferotemporal cortex that familiarity training leads to the sparsification of population neural representation to the familiar stimuli, as neurons’ responses to their preferred familiar stimuli were found to be enhanced, while their responses to non-preferred familiar stimuli were suppressed, resulting in a sharpening of the stimulus selectivity tuning curves of the neurons [6, 7].

In the early visual cortex, Huang et al. [8] showed that neurons with localized receptive fields became sensitive to the global context of familiar images. Based on timing, it can be inferred that this sensitivity is mediated by the recurrent circuits within V2 rather than feedback from higher visual areas. Similar effects have also been observed in V1 as well, but with a shorter delay with stimulus onset, significantly earlier than the familiarity effects in IT. These observations suggest a rapid plasticity mechanism in the early visual cortex modifying the recurrent circuit within each visual area along the visual hierarchical system to encode global or semi-global familiar image context. These findings, together with the involvement of BCM-like learning rule in the familiarity effect [9, 10], suggest that neurons in the early visual cortex, with local receptive fields, can rapidly learn recurrent excitatory circuits to encode global images.

However, the computational rationales and the neural mechanisms of this rapid plasticity are not well understood. Proposals on the behavioral and perceptual benefits of the familiarity effects in IT have been focused on image discrimination, reduced saliency to familiar entities, and novelty detection [6, 7, 11, 12]. Recent studies found that representations of natural movies in the primary visual cortex change with repeated exposures [13], which inspires us to investigate the relation between representational changes and familiarity training. The objective of this study is to link the familiarity effects to manifold learning, which seeks to find the correct geometric relations between visual concepts and their variants generated by different smooth transformations, such as different levels of occlusion, view angle, etc. [14, 15, 16, 17] (here the term “visual concepts” refers to images with different semantic meanings, in contrast to the general term “images” that refers to both visual concepts and their variants).

In this paper, we develop a V1-based neural circuit model based on BCM Hebbian learning and other standard V1 circuitry elements that can account for the familiarity training effects. This is a universal canonical circuit motif that can be generalized to V2, V4, and IT. We analyzed this circuit to show that familiarity training of a certain global image stimulus, henceforth called a global visual concept, transforms the neural representation manifold in such a way that unimportant variations of the same concept are ignored while discrimination of different visual concepts is maintained. We demonstrated that this manifold transform provides a more robust encoding of familiar images or concepts so that they are more immune to noises. We found that the recurrent cortical circuit implements the manifold transform by locally linear computations that rotate the slow patterns of the local network dynamic to a direction in the input space that is orthogonal to the variation of the same concept. This novel perspective on cortical recurrent circuits provides insights into the functional rationales underlying the familiarity learning observed in the visual cortex.

## 2 Results

### 2.1 Recurrent neural circuit model of visual cortex shaped by BCM learning

Familiarity training effects have been reported in macaque ITC and V2. Similar effects have also been observed by us in macaque V1 (unpublished observations), as well as by others in mouse V1 [12, 18, 19]. Since V1 circuits are relatively well understood and modeled, we decided to use the V1 model for this study. We constructed a neural circuit model of the primary visual cortex to demonstrate that the plastic horizontal connections, driven by the BCM rule [9, 20, 21], can reproduce familiarity effects. This basic associative learning mechanism is likely generalizable to model familiarity effects in other visual and non-visual cortical areas.

The network model (Fig 1A) is a firing-rate-based recurrent neural network with *N*_*h*_ hypercolumns (with *N*_*r*_ rows and *N*_*c*_ columns). Each hypercolumn comprises *N*_*d*_ excitatory neurons with receptive fields (RF) derived from sparse coding [22, 23]. We have *N*_*e*_ = *N*_*r*_ × *N*_*c*_ × *N*_*d*_ excitatory neurons and the same number (*N*_*i*_) of inhibitory neurons in the network. Each excitatory neuron *k* receives projection from its excitatory neighborhood (*NE*(*k*)) with a spatial extend of *R*_*e*_ (Fig 1B). Each inhibitory neuron *k* receives projections from the excitatory neurons of the same feature channel located in its inhibitory neighborhood (*NI*(*k*)) with range *R*_*i*_ (Fig 1B) and projects back to all excitatory neurons in the network, mediating iso-orientation (or iso-feature) suppression. For neurons not at the border, |*NE*(*k*) | = *N*_*d*_ × (2*R*_*e*_ + 1)^2^, and |*NI*(*k*)| = (2*R*_*i*_ + 1)^2^. In addition, excitatory neurons within the same hypercolumn project to an inhibitory neuron, which uniformly inhibits these excitatory neurons in return, as a form of divisive normalization [24, 25].

**Fig 1.**
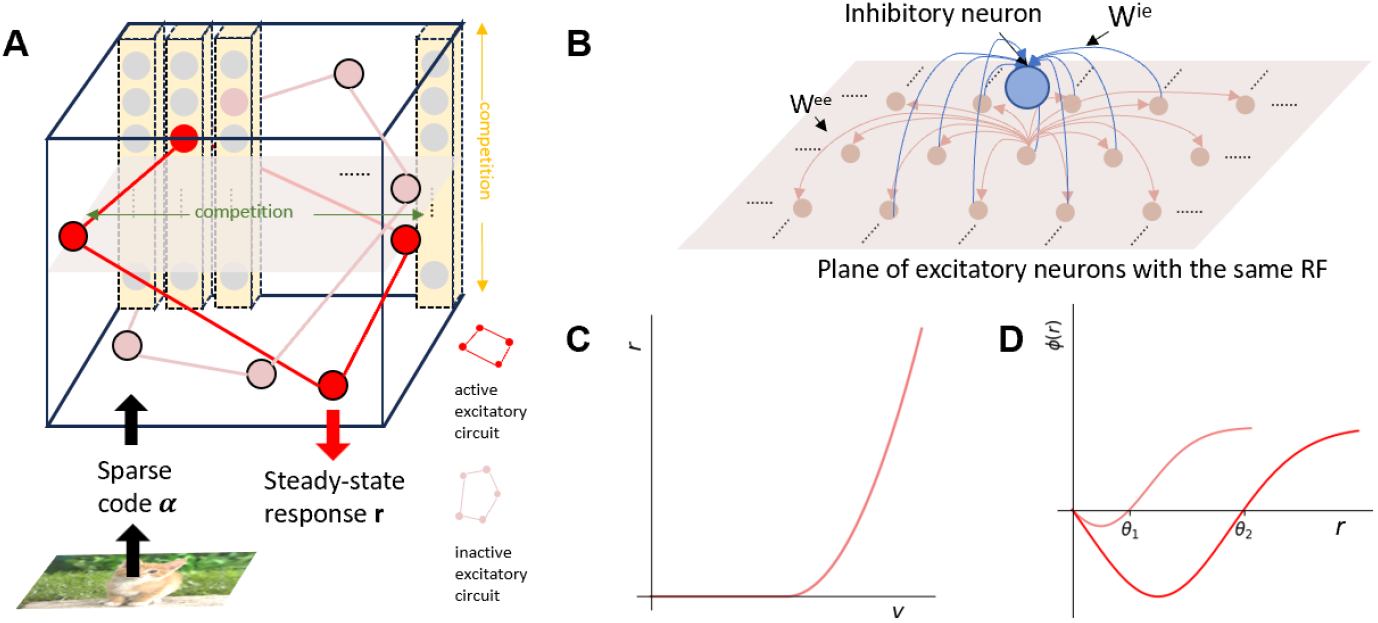
Recurrent circuit model of the visual cortex. (A) Illustration of the network structure. An excitatory sub-circuit is formed between excitatory neurons across different hypercolumns for encoding an image. Competitions exist among neurons within the same feature channel across hypercolumns and among neurons of different feature channels in the same hypercolumn. The input to the network or the neurons’ initial responses ***α*** are computed as the sparse codes of the image [22], and the output ***r*** is the steady-state population response of the excitatory-inhibitory network. (B) Illustration of the excitatory neighborhood (for a single feature channel) and inhibitory neighborhood. (C) The activation function of a neuron that maps membrane potential *v* to neuronal firing rate *r*. (D) The graph illustrates the BCM learning rule, showing the synaptic potentiation *ϕ* of the post-synaptic neuron depends on its firing rate and the BCM threshold. The synaptic weight increases when the firing rate is above the BCM threshold and decreases otherwise. BCM rules for two thresholds *θ*_1_,*θ*_2_ are shown.

The dynamics of the excitatory population and inhibitory population are given, respectively as:

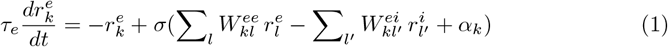

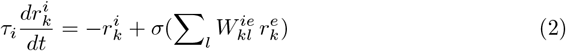

where 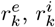 are the firing rates of the *k*^*th*^ excitatory neuron and inhibitory neuron, respectively. 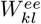 is the E-E connection from excitatory neuron *l* to excitatory neuron *k*; similarly 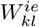 and 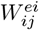 are the E-I and I-E connection; *α*_*k*_ is the input to the excitatory neuron *k* obtained via convolutional sparse coding [23]. 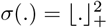 is the squared relu activation function (Fig 1C), as introduced in [25]. *τ*_*e*_ and *τ*_*i*_ are the time constants of excitatory and inhibitory neurons, respectively.

Here, we are pursuing a minimal circuit mechanism that could reproduce the familiarity effect in neural circuits. Hence, we start by only considering excitatory plasticity while assuming that familiarity learning did not fundamentally change the canonical inhibitory mechanisms. Hence, only *W* ^*ee*^ is subject to BCM learning, while *W* ^*ei*^ and *W* ^*ie*^ are fixed. The initial value of the E-E connection is set to *w*_*ee*_*/* |*NE*(*k*) |, within the excitatory neighborhood. For a single inhibitory neuron, the E to I connection to this neuron is uniformly set to a fixed value *w*_*ie*_*/* |*NI*(*k*) | within its inhibitory neighbors. The I to E connection from this inhibitory neuron to all excitatory neurons is set to the fixed value 1*/N*_*i*_. *w*_*ie*_ is set so that normalization within the hypercolumn and iso-orientation (iso-feature) surround suppression across the hypercolumns are strong enough to ensure the stability of the network.

The BCM learning rule [21] is written as:

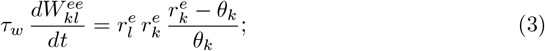

where *τ*_*w*_ is the synaptic time scale determining the speed of learning. The BCM threshold *θ*_*k*_ for excitatory neuron *k* is computed by taking the exponential moving average of the neuron’s squared firing rate. The synaptic connection strength increases only if the neuronal firing rate exceeds the BCM threshold, and decreases if the firing rate is below the threshold. As the threshold traces the neuron’s activity, for neurons after a period of high activity, the threshold increases (red curve in Fig 1D), and for neurons after a period of low activity, the threshold decreases (pink curve in Fig 1D). This negative feedback mechanism prevents unbounded synaptic increase and ensures the stability of the learning. We also added synaptic scaling to preserve the total strength of pre-synaptic connections to an excitatory neuron *i* throughout the learning [26]:

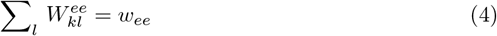

### 2.2 Familiarity suppression and tuning curve sharpening in the model

We verified that the model could indeed reproduce the effect of familiarity suppression. Fig 2B shows the averaged peri-stimulus firing rates of all excitatory neurons across a set of stimuli before and after familiarity training. Neural activity in the network peaks right after stimulus presentation. Surround suppression kicks in, leading to the response decay. This is followed by some small oscillations due to the interaction of the surrounding excitatory and inhibitory neurons. Finally, the activity settles into a steady state. After familiarity training, the averaged population responses to trained stimuli are profoundly suppressed (Fig 2B, yellow area), consistent with the findings of the neurophysiological experiment in the macaque visual cortex (Fig 2A).

**Fig 2.**
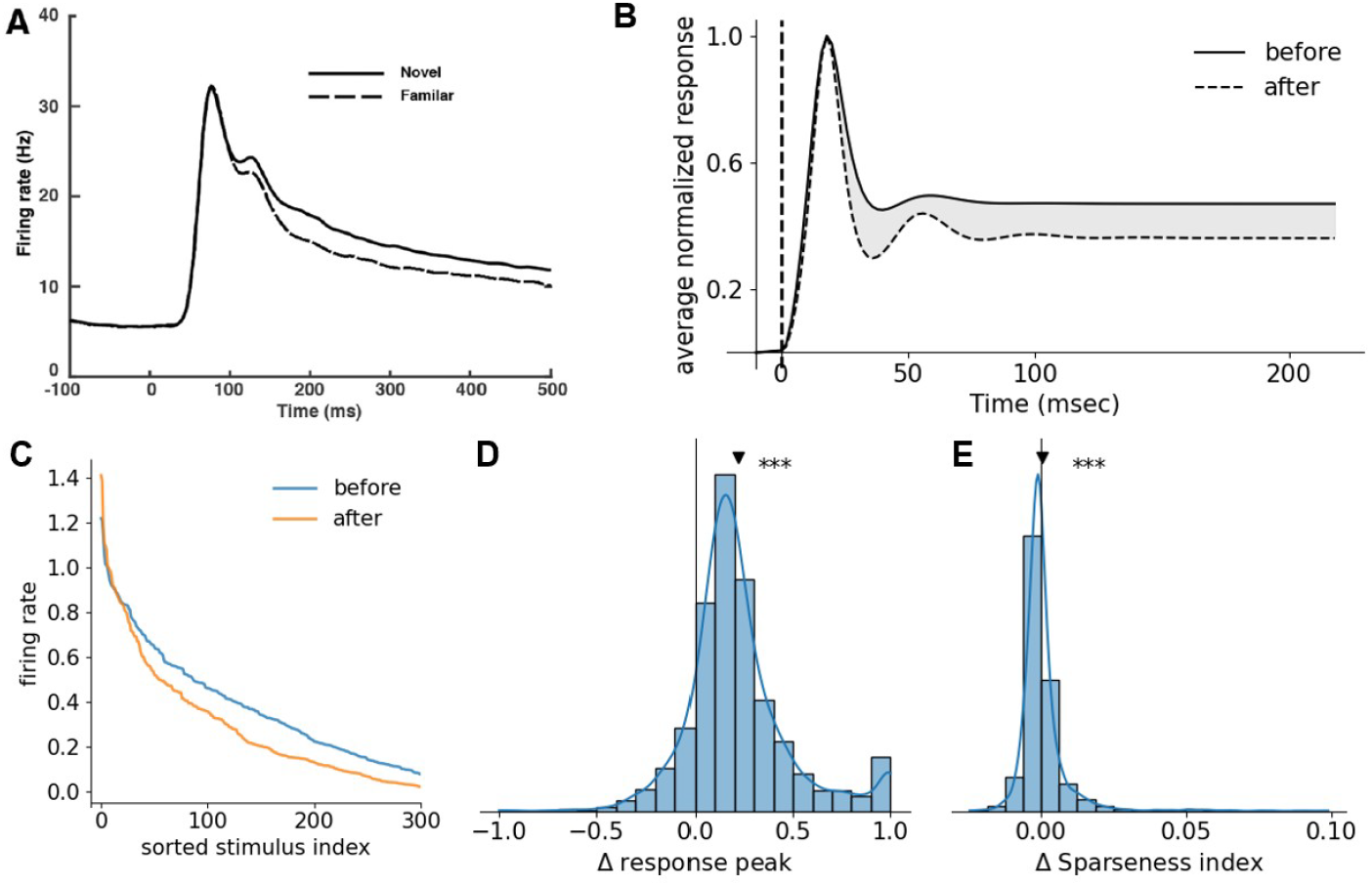
Tuning curve sharpening and familiarity suppression. (A) Experimental observation showing neural responses of familiar images are suppressed relative to novel images (Huang et al. 2018). (B) The average neural population response to a set of images is suppressed after the neurons are subjected to familiarity training. The grey area represents the suppression caused by familiarity training. This is equivalent to comparing the neural population responses to a set of trained images against a set of untrained novel images in the neurophysiological experiment, in which neurons could not be tracked and compared reliably across days. (C) Example neuron’s tuning curve to 500 images before and after familiarity training. After training, the tuning curve becomes sharper: the response to the most preferred familiarized stimulus is enhanced, and responses to non-preferred familiarized stimuli are suppressed for this neuron, resulting in increased sparsity of the neuronal tuning. (D) The population histogram of change in the tuning peak shows a significant increase. (E) The population histogram of change in sparsity shows a small but statistically significant increase due to the long right tail. *** indicates *p <* 0.05 for the alternative hypothesis that the mean is greater than 0.

The trained network exhibited significant sharpening of the stimulus tuning curves of the excitatory neurons to the familiar stimuli. Fig 2C shows sorted tuning curves of an example neuron before and after training, where the index was ordered by pre-trained response (in descending order). Familiarity training suppressed its response to most stimuli while only increasing the response to the strongly preferred stimuli, resulting in a sharper tuning curve. We quantified the relative change in the sparseness index (see Section 4.3) of the stimulus tuning curves for all neurons by the ratio ((after - before) / (after + before)). The overall effect on the tuning sparseness across the population is small but still significant, attributing to the long tail right tail of the distribution (Fig 2E), where about 3% of neurons show large changes (2.5%-10%) in the tuning sparseness before and after training. Meanwhile, the peak firing rates of individual neurons are increased significantly (Fig 2D). These results suggest that the familiarity suppression effect can result from the sparsification of the neural representation in response to familiar stimuli. The increase in tuning peak reflects the formation of a local circuit based on Hebbian learning to encode a particular global stimulus. When a familiar stimulus is presented, the neurons involved in this local circuit will boost each other’s response, leading to an increase in the firing rates of neurons involved in the circuit. On the other hand, the enhanced responses of neurons involved in the circuit will suppress all the other neurons not involved in the circuit in that neighborhood due to the surround suppression and within-hypercolumn normalization, resulting in a net decrease in the averaged responses in the general population.

The connectivity strength in the spatial cortical network is typically modeled distance-dependently [25, 27, 28, 29], while for simplicity the fixed inhibition strength in our network is uniformly distributed in space. We experimented a form of spatial dependent inhibition *w*_*ie*_ following a Gaussian profile (up to *σ* = 1) (Fig S1A) but found them inconsequential to familiarity suppression and the sharpening of tuning curves (Fig S1B-C). Having strong enough inhibition, however, is crucial. We varied *w*_*ie*_ to assess the impact of inhibition strength. We found that the network would be unstable when the inhibition strength is too weak. However, the main effects (familiarity suppression and tuning curve sharpening) can be observed as long as the network is in a stable regime, with the effect decreasing as *w*_*ie*_ moves away from the critical value where bifurcation happens (Fig S2A-B).

### 2.3 Manifold transform in the recurrent cortical circuit

In this section and the following, we investigate the computational rationales underlying the learning of the circuit associated with the familiarity effects. Computationally, our model is an attractor network; the input to the network is the sparse code ***α***, and the network output is the steady-state response ***r***, i.e., the representation of the stimuli. A neural representation manifold is defined as a set of representations of a set of stimuli. [30, 31]. The recurrent circuit performs a manifold transform operation that is dynamically modulated through learning, which maps the input representation manifold to the output representation manifold. We are interested in how learning modulates the manifold transform and what computational benefit familiarity training could bring.

Inspired by the recently proposed sparse manifold transform framework [16, 32], we hypothesize that the manifold transform mediated by the recurrent network would increase the robustness of the representation of visual concepts, which compensates for the low reliability of the sparse code. In the following paragraph, We first discuss the problem of the neural representation caused by the sparse coding algorithm, and how the manifold transform makes the neural representation manifold better reflect the true geometry of the natural images manifold. Then, we propose a more general manifold transform that can be implemented by recurrent cortical circuits.

#### The objective of manifold transform

Studies on manifold learning [14, 16, 33] have long noticed that the primary sensory areas, such as V1, with neurons selective to local features, such as the sparse code dictionary used in our model, cannot represent the geometry inherited in images. Smooth transitions in images are often represented by abrupt changes in the population neural codes, particularly when they are sparse codes. Consider the smooth manifold in the high-dimensional space of local visual features (Fig 3A), as discussed in [16], the learned spare code dictionaries are discrete samples on the manifold ℳ, and the sparse code ***α***_*i*_ of an image stimulus ***x***_*i*_ is a composition of multiple delta functions on the manifold domain (Fig 3A). The underlying transformation of the stimulus, such as those induced by various degrees of noise and occlusion, or the changing view angle, is a collection of flows in ℳ . Because the learned dictionaries are typically unordered, the flow on the manifold leads to a drastic change in the sparse code (Fig 3A).

**Fig 3.**
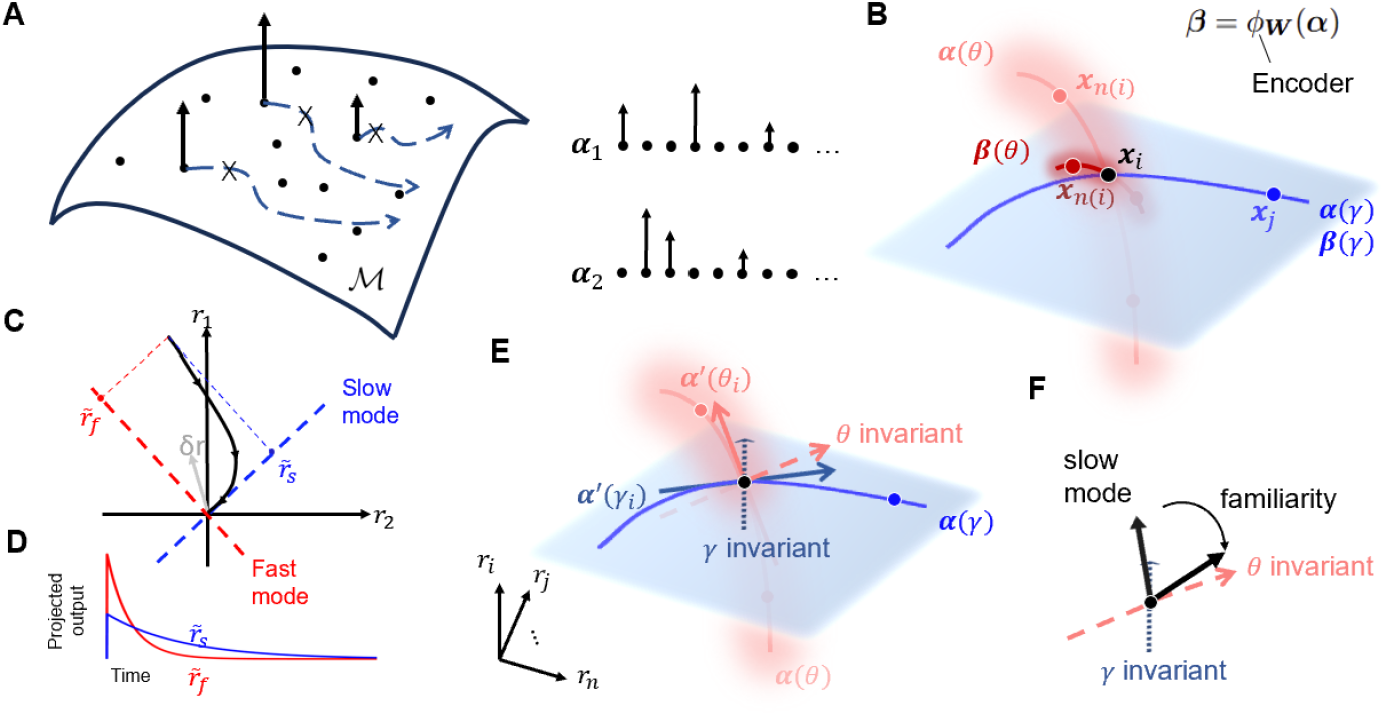
Manifold transform in the recurrent cortical circuit. (A) A conceptual model demonstrating how sparse code disrupts inherited geometric structures in images. ℳ is the putative feature manifold; black dots represent sparse dictionaries that are landmarks on the manifold; black arrows are delta functions on the manifold domains constituting the sparse code of an image; blue arrows show the direction of flows; ***α***_1_, ***α***_2_ are two example sparse codes of two images along the flow. (B) Illustration of manifold transform. The blue area represents the smooth manifold of global concepts (concept manifold), and the blue curve shows a function on the concept manifold, determined by parameter *γ*, where stimulus ***x***_*i*_ (black dot) smoothly changed to the non-neighbor stimulus ***x***_*j*_ (blue dot) in the representation space. The red area represents the manifold of diverse variations of the concept (variants manifold), and the red curve is a function on the manifold determined by parameter *θ*, along which ***x***_*i*_ changes to the neighbor stimulus ***x***_*n*(*i*)_ (red dots) in the representation space. Assuming no changes in the concept manifold (for visual simplicity), the learned encoder *ϕ*_***W***_ compresses the variants manifold. (C) Illustration of dynamical mode decomposition. The black curve is the decaying trajectory of the neuronal response to the small perturbation *δr* around the fixed point (origin). The red and blue dashed line denotes fast and slow dynamical modes, respectively. 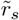 and 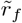 indicate projected neuronal response onto the slow and fast modes, respectively. (D) shows the temporal evolution of 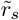 and 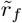,which are disentangled, exponential decay with fast decaying speeds (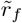,red curve) and slow decaying speed (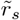,slow curve). (E) Illustration of *θ* invariant direction and *γ* invariant direction. ***α***^*′*^(*θ*_*i*_): derivative of ***α*** w.r.t. *θ*; ***α***^*′*^(*γ*): derivative of ***α*** w.r.t. *γ. θ* invariant direction and *γ* invariant direction are orthogonal to ***α***^*′*^(*θ*_*i*_) and ***α***^*′*^(*γ*). (F) Training should render slow modes of the network dynamics more aligned to *θ* invariant direction than to *γ* invariant direction, leading to neural responses that are more similar to the neighbor stimuli than other stimuli.

Here, we consider the problem setup where the image stimuli ***x*** comprise visual concepts ***c*** as well as their variants generated by the smooth transformation, denoted by {***c***^*′*^}. We define the variants manifold associated with a concept to be 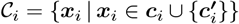, and the concept manifold to be a smooth manifold underlying all the visual concepts. Our goal is to find an encoder *ϕ*_***W***_, parameterized by ***W***, that takes sparse code ***α***_*i*_ as input and captures the true geometric relation between stimuli in the manifold of its representation (denoted by ***β***). To do that, the learned representation should satisfy the following property compared to the sparse code: Locally around each stimulus ***x***_*i*_ ∈ 𝒞_*i*_, ***x***_*i*_ should be closer to neighbor stimuli ***x***_*n*(*i*)_ that are in the same variants manifold 𝒞_*i*_ relative to stimuli ***x***_*j*_ that are in other variants manifolds 𝒞_*j*_. In other words, the manifold transform should compress all the variants manifolds relative to the concept manifold (Fig 3B).

The desired representation can be obtained by maximizing the similarity of representations within the variants manifolds of each concept while maintaining the difference between the representations of the different concepts. Various objectives have been proposed in the manifold learning and self-supervised learning literature to address this trade-off [14, 16, 17, 34, 35, 36]. Here, we considered an objective that maximizes the variant similarity with a soft constraint by optimizing the parameter ***W*** of the encoder.

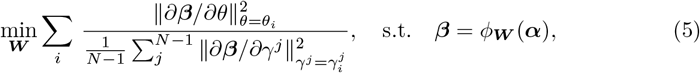

The objective operates on the derivatives of the *θ* function and the *γ*_*j*_ function on the variants manifold and concept manifold, respectively. The former controls the smooth change from a concept ***x***_*i*_ to its neighborhood stimuli ***x***_*n*(*i*)_ in a variants manifold, and the latter controls the smooth change from a concept ***x***_*i*_ to other concepts ***x***_*j*_ (Fig 3B). This objective seeks to minimize the change of ***x***_*i*_ to ***x***_*n*(*i*)_ relative to the averaged change of ***x***_*i*_ to different ***x***_*j*_, thereby ignoring the variations within the same concept while preserving the variation between the distinct concepts.

#### Locally linear manifold transform by the recurrent cortical circuit

The traditional manifold transform algorithms [15, 16] typically use a single linear transform for all the visual concepts. However, organizing the representational geometry of all the concepts and their variations with this linear approach could be challenging. A more flexible way might be to let the manifold transform manipulate the local variations around different points on the manifold with distinct linear operations, thereby enabling concept-specific transforms while maintaining global nonlinearity. The recurrent circuit is inherently suitable for accomplishing this type of globally-nonlinear and locally-linear manifold transform. The nonlinear dynamic of a recurrent circuit can be approximated by a linear dynamical system determined by a Jacobian matrix (see Section 4.4) locally around each fixed point [37, 38, 39]. As a result of this property, the manifold transform performed by the recurrent cortical circuit can be seen as a composition of multiple locally linear transforms around each stimulus in the representation space. Therefore, the recurrent circuit could potentially offer greater flexibility to the manifold transform.

Here, we consider *ϕ*_***W***_ to be the recurrent cortical circuit, and its steady-state response ***r*** to be the output of manifold transform (***β*** in Eq 5). To see how the recurrent circuit performs locally linear manifold transform, we consider the tuning derivative defined as ***r***^*′*^ = *∂****r****/∂θ*. The tuning derivative evaluated at *θ*_*i*_ can be decoupled into disentangled components that fall along each dynamical mode, obtained by eigendecomposition of the Jacobian matrix of that fixed point (see Section 4.4). The dynamical modes are a new set of bases along which the projected local dynamic around a fixed point is decoupled into independent, exponential evolution with different time scales (Fig 3C, D). The basis with a shorter time scale (faster evolution) is the fast mode, whereas the basis with a longer time scale (slower evolution) is called the slow mode. The decomposition of tuning derivative at *θ*_*i*_ w.r.t. dynamical modes is written as

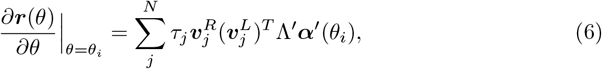

where *τ*_*j*_ represents the dynamical timescale of the *j*^*th*^ mode, derived from the eigenvalue of the Jacobian: *τ*_*j*_ = −1*/Re*(*λ*_*j*_). The vector 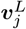 denotes the *j*^*th*^ dynamical mode, which is also the left eigenvector of the Jacobian. The vector 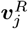 is the corresponding right eigenvector. Λ^*′*^ is the sensitivity matrix (see Section 4.4), and ***α***^*′*^(*θ*_*i*_) is the input derivatives at *θ*_*i*_. Eq 6 shows that the derivative along the representation manifold is the superposition of input derivatives processed by different modes, weighted by their respective time scales. The dynamical mode 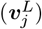 filters the input derivatives by projection, and the filtered information is projected back to the neural space using right eigenvectors 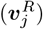 as the basis. Therefore, the filtering performed by the dynamical modes with slow time scales (large *τ*) determines what information in the input would be extracted or discarded.

Taking advantage of the orthogonality between left and right eigenvectors, a simpler linear relationship between input ***α***(*θ*) and output ***r*** can be obtained by projecting the representation onto the slow dynamical modes:

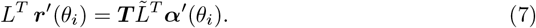

The column of *L* is 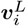 and the column of 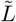 is 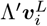.The matrix ***T*** = *δ*_*ij*_*τ*_*i*_, containing the time scales of each mode in the diagonal. Eq 7 shows that the slow dynamical modes constitute an efficient linear readout for the geometry of the representation manifold, as the representation projection onto each slow mode could retain the geometry-relevant information from the corresponding input projection through linear scaling. In contrast, the arbitrary projection of the representation is the superposition of input projections onto different dynamical modes, thus mixing the geometry-relevant and -irrelevant information.

The key aspect of the local linear transform lies in how slow modes filter inputs. We define the *θ*-invariant direction to be orthogonal to the input derivatives ***α***^*′*^(*θ*_*i*_), along which change in *θ* is accompanied by little change in neural input activity ***α***.

*γ*-invariant direction is defined similarly with respect to change along the concept dimensions *γ*. (Fig 3E). Eq 6 and Eq 7 imply that the alignment of the network’s slow modes with the invariant direction modulates the similarity of the stimulus representations. Minimization of the objective (Eq 5) requires the slow modes to align more to the *θ*-invariant directions than the *γ*^*j*^-invariant directions (Fig 3F), so that the representation ***β*** changes more slowly w.r.t. *θ* than w.r.t *γ*^*j*^ around each fixed point. Consequently, the manifold’s global geometry is reshaped through local optimization of stimulus representation similarities.

### 2.4 Simulation to relate familiarity-trained recurrent cortical circuit and manifold transform

Manifold transform, achieved by optimizing the objective function in Eq 5, by design, seeks to compress the variants manifold, i.e., manifold formed by variations of a visual concept, relative to the concept manifold, i.e., the smooth manifold underlying all visual concepts. The variation of a visual concept can come in a variety of forms, such as noises and occlusions or changes in images associated with views. Manifold transform seeks to ignore these variations within a concept while maintaining sensitivity to the distinction between the different concepts.

Would familiarity training induce the same effect on a recurrent circuit, compressing the irrelevant variations such as noises and occlusion for each visual concept? To test this prediction, we perform a simulated neurophysiological experiment on spatial and temporal associative learning. The experimental paradigm is as follows: the network, simulating a monkey performing a fixation task, will be presented with a pair of stimuli.

The first image is a noise-corrupted version of the second image called the target image. The network is trained for five epochs. Each epoch involves 150 steps of BCM learning (Fig 4A). The noises are salt and pepper occlusion noises at four levels: 0%, 10%, 30%, and 50% (Fig 4A). Corruption of target image *l* with noise level *n* results in a conditional distribution of stimuli *p*(***x*** | *n, l*). We draw 10 image samples denoted by ***x***_*n*,*l*_ from each conditional distribution. Here, the visual concepts are the target images, and the noise level for each image concept controls the variations of the input for that image concept, parameterized by the *θ* in the previous section.

**Fig 4.**
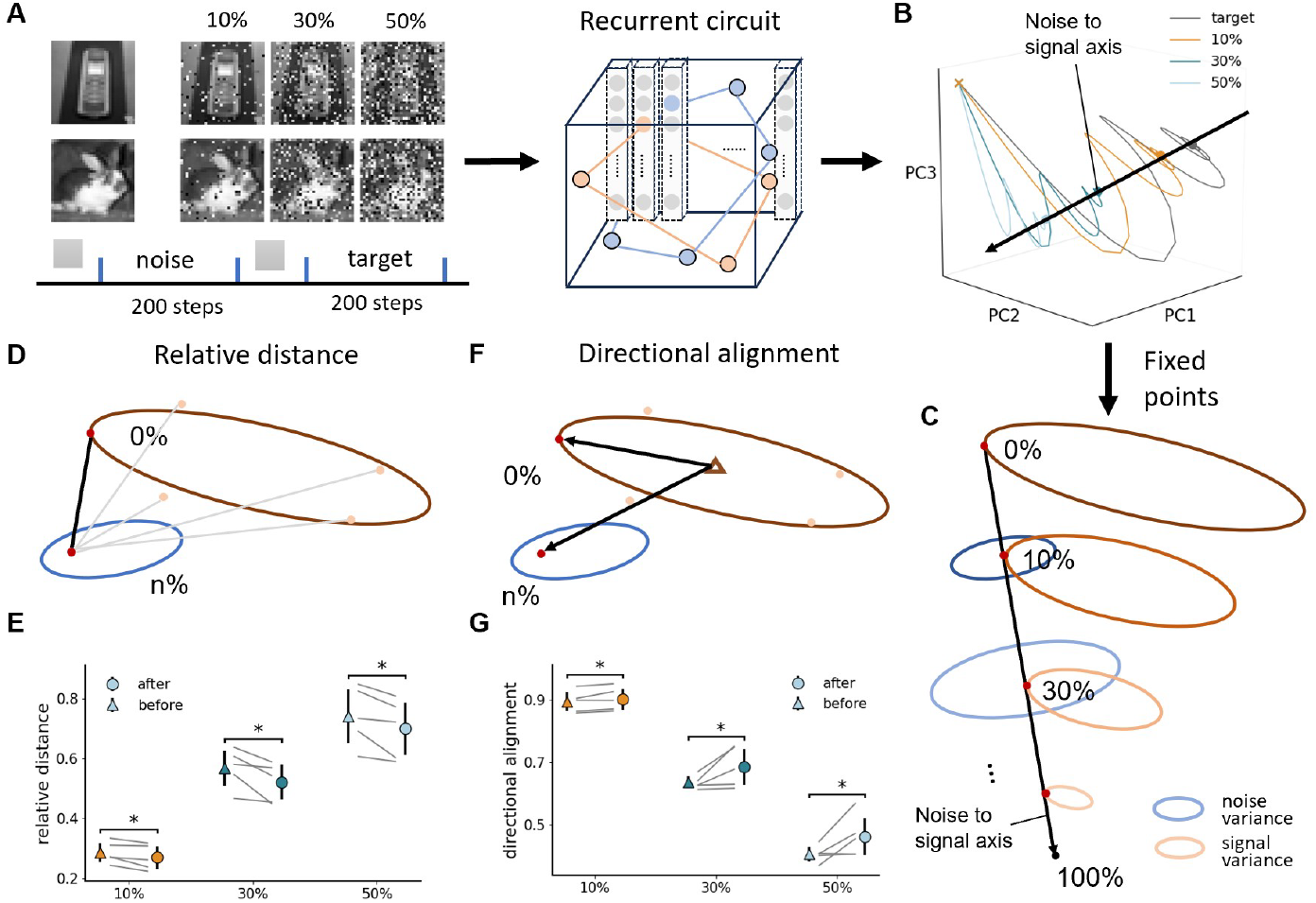
Manifold transformation in the familiarity association experiment. (A) Upper: Example images corrupted with different levels of noise. Lower: familiarity training for one noisy-target image pair. (B) Trajectories of different noise levels correspond to an example image in the model. The trajectory is averaged across noisy image samples. The black arrow indicates the direction along which the noise level changes (denoted as the image-to-noise axis). Cross: trial start; Dots: trial end. (C) Cartoon illustration of the cone formed by decreasing signal variance and the cone formed by increasing noise variance of an image. (D) Illustration of the relative distance. The black line between two red dots represents the Euclidean distance between the noisy image and the target image, which is normalized by the sum of distances from this noisy image to all other clear images (represented by the gray lines) (E) Relative distance significantly decreases after training for all noise levels. Triangle marker: before training, circle marker: after training (epoch5 for the simulation). Grey straight lines are for individual target images. *: *p <* 0.05. Error bar represents std across target images. (F) Illustration of the directional alignment. The triangle denotes the reference point formed by averaging all the target images. Cosine similarity is calculated between the two vectors pointing from the reference point to the noisy image and its target. (G), Similar to E but for the directional alignment, which significantly increases after training for all noise levels

Familiarity training in this experiment is designed to induce temporal association of images and spatial association of visual features within each image. Given the noisy image is followed by a clear version of the image, this is a form of temporal association learning between the input images, reminiscent of Slow Feature Analysis [33]. The repeated presentation of the target image during training will strengthen the connection between the neurons coding the spatially occurring events in a familiar image based on the Hebbian learning mechanism. If familiarity learning in a recurrent network is consistent with the manifold transform behaviorally, the learned representation in the recurrent circuit should ignore the variations induced by noise (i.e., the *θ* dimension) and maintain the discriminability among the different target images (i.e., the *γ* dimension) in the manifold, resulting in greater robustness against the variation due to noise.

After each epoch of training, we probed the neural activity and analyzed the neural representation manifold formed by fixed points of the excitatory population, denoted by ***r***^***e***^. The familiarity suppression and sharpening of stimulus tuning and population tuning are also observed in the familiarity association experiment (Fig S3A-C). The evolution of the population activity of excitatory neurons in response to different noise levels forms distinct trajectories in the neural representation space, which start at the same initial resting state of the network, then diverge and settle into different fixed points corresponding to the distinct input images (Fig 4B). Fixed points of different noise levels are organized along certain directions in the neural representation space (Fig 4B), forming an image-to-noise axis along which the noise level changes gradually. We will show that familiarity training would compress this axis to increase the noise-robustness of the representation.

Next, we investigated the geometry of the neural representation manifold. We found that the neural representation lies on a lower dimensional space as compared to that of the input sparse code (Fig S3G). The neural representation manifold formed by all the target-noise image pairs has approximately a cone shape, as shown in figure 4C. Specifically, The 10 samples from each conditional distribution of stimuli given the noise level and visual concept form different clusters. The target image is located at one end of the manifold, and increasing the noise level drives the cluster mean farther away from the target image, accompanied by an increase in the variance of the distribution (noise variance)(Fig S3D). Such a geometry arises because the higher noise level reduces the similarity between different noisy image samples. In the meantime, as the noise level increased, the distances between cluster means in the same noise level (signal variance) decreased. For 100% noises, all the “images” will converge to the same cluster, reflecting the reduced signal contents in the corrupted images. Therefore, if only the cluster means are considered, the manifold’s shape is also a cone but has the opposite direction compared to the first cone (Fig S3E).

We then quantitatively measured whether the trained network transformed the representation manifold in a way consistent with the manifold transform specified in Eq 5. We defined the relative distance as a discrete approximation of the objective (Eq 5). The relative distance between the neural representation of a noisy image and the target image is their Euclidean distance, normalized by the sum of the distances between that specific noisy image and all the clear images except its target (Fig 4D). The directional alignment is another metric to measure the noise robustness, which is the cosine similarity between the vector from a reference point to the representation of the noisy images and the vector from the reference point to the representation of the target image, where the reference point is the mean of all the target image representations (Fig 4F). We found that for all noise levels, the directional alignment increases and the relative distance decreases, comparing before and after training (Fig 4E, G). Consistent with this compression effect, we observed a decrease in the dimensionality of representation after training (Fig S3G). These results show that the familiarity-trained network behaves in a way consistent with the manifold transform, ignoring the variation specifically arising from the noises, thereby enhancing the noise-robustness of natural image representations.

### 2.5 Familiarity training learns manifold transform by aligning the slow modes of network dynamics to the *θ*-invariant dimension

We performed numerical experiments to evaluate the predictions of our analysis on the locally linear manifold transform induced in the recurrent circuit by familiarity training. We used numerical methods to perform linearization analysis around each fixed point of the network. The dynamical modes around each fixed point are all decaying, suggesting that all the fixed points are stable attractors. The decaying modes can be categorized into several groups according to the decaying time constants. The first group of decaying modes has time constants greater and gradually approaches *τ*_*e*_ (the time constant of excitatory neurons, see Eq 1); The second group has *τ* = *τ*_*e*_, but the modes are all zero vector, thus will not contribute to the dynamic; The third group has time constants smaller than *τ*_*e*_ and larger than *τ*_*i*_ (the time constant of inhibitory neurons, see Eq 2), and the last two groups have *τ* = *τ*_*i*_, decaying with the same speed as the inhibitory neurons (Fig 5A). Eigenvectors in groups 1 and 3 have exclusive selectivity for excitatory neurons. Hence, they are relevant to the manifold formed by excitatory population responses. In contrast, eigenvectors in group 4 constitute mainly inhibitory loading (Fig 5B). These observations suggest that the group 1 dynamical mode is most relevant to the transformation of the neural manifold formed by excitatory population response.

**Fig 5.**
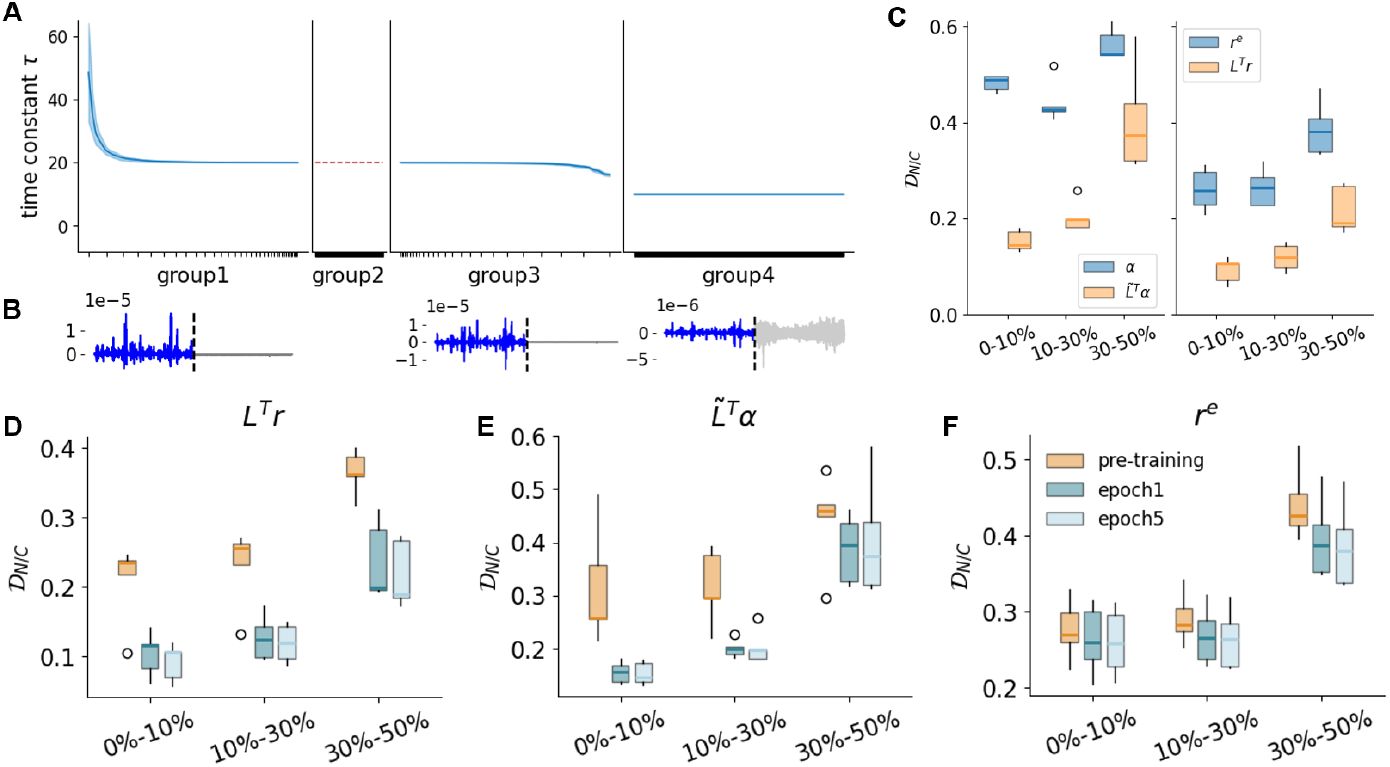
Familiarity training aligns slow dynamical modes with *θ*-invariant direction. (A) Groups of dynamical modes that are specified by decaying the time constant *τ* . First panel: Group 1, with *τ > τ*_*e*_; Second panel: Group2, with *τ* = *τ*_*e*_ and all eigenvectors are zeros vectors. Third panel: Group 3, with *τ*_*i*_ *< τ < τ*_*e*_. Last panel: Group 4 with *τ* = *τ*_*i*_. The blue line represents the mean, and the shaded area represents s.e.m. across visual concepts, noise levels, and training epochs. (B) Averaged eigenvectors of groups 1, 2, and 4 correspond to excitatory neurons (colored in blue) and inhibitory neurons (colored in gray). Eigenvectors in group 1 and group 3 are irrelevant to inhibitory neurons, while eigenvectors in group 4 primarily represent loadings to inhibitory neurons. (C) Normalized noise distance 𝒟_*N/C*_ in the epoch5 trained network decreases in the projection space. Left panel: comparing feedforward input (***α***) and input projection 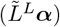;Right panel: comparing the original representation (***r***^***e***^) and response projection (*L*^*T*^ ***r***). Boxplots show distributions of visual concepts. (D-F) Familiarity training continuously reduces normalized noise distance 𝒟_*N/C*_ for input projection, response projection, and original neural response. Boxplots show distributions of visual concepts.

In this experiment, target images represent distinct visual concepts, where variations within each concept are governed by the noise level, parameterized by *θ*. We refer to the *θ*-invariant direction as the direction insensitive to noise variation, whereas *γ*^*j*^-invariant direction as the direction insensitive to the changes in concepts. The alignment of the top 100 slow modes of the trained network with noise-invariant directions and concept-invariant directions is examined. We used the distance between two adjacent noise levels (noise distance 𝒟_*N*_) and the distance between concepts (concept distance 𝒟_*C*_) to approximate the two tuning derivatives (see Section 4.7 and Fig S4A). We found that the slow decaying dynamic modes aligned more with the noise-invariant direction than the concept-invariant directions, evidenced by the decreased normalized noise distances (ratio of noise and concept distance, 𝒟_*N/C*_) in the projected input 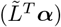 compared to the inputs (***α***) (Fig 5C). Projecting response onto the slow modes (*L*^*T*^ ***r***) further decreases the normalized noise distance (Fig 5C), confirming that slow modes constitute an efficient subspace for the noise invariance.

We then investigated the effect of training by comparing the trained network with the pre-trained network. For subspace spanned by the top 100 slow modes, training decreases the normalized noise distance in input projection and response projection around each fixed point (Fig 5D-E). The results suggest that familiarity training facilitates manifold transformation by continuously improving the local alignment of slow dynamical modes with noise-invariant direction, making the slow subspace more robust against noise. We also observed the decrease in the normalized noise distance in the original neural representations (***r***^***e***^) (Fig 5F), which is consistent with the finding of relative distances in Fig 4E.

Furthermore, we asked whether the observed effects are consistent across different project subspace dimensions, i.e., the number of slow modes we used to construct the projection subspaces. We found that the first group of decaying mode consistently aligned with the noise-invariant directions more so than the concept-invariant direction (Fig S4D), and noise invariance in the slow modes subspace spanned by different numbers of group 1 modes is consistently better than the original response (Fig S4E). The effects of familiarity training are presented in a wide range of subspace dimensions. We quantified the training effect by the Pearson correlation coefficient between training epochs and normalized noise distance. Fig S4F-G shows that the training effect of the input and output projection exhibits a decreasing trend, with saturation at higher subspace dimensions. Therefore, the observed effects are consistent with different projection subspace dimensions. The result also emphasizes the critical role of the first quarter of the slow modes in compressing the irrelevant variation, as further adding more slow modes yields diminishing effects.

Taken together, these results demonstrate that recurrent networks align the slow dynamical mode specifically to the *θ*-invariant direction, and familiarity training continuously enhances the alignment. These properties of the network dynamics facilitate the formation of a representation manifold geometry that improves the representation of familiar stimuli and their robustness against noise.

## 3 Discussion

In this work, we showed that a neural circuit model with standard V1 motifs of hypercolumns, local sparse codes, and local inhibitory and excitatory mechanisms can rapidly learn to encode global image concepts by modifying the excitatory connections in the near-surround using a BCM Hebbian learning mechanism. BCM rule has been used to explain the development of the receptive field’s orientation selectivity [21]. The BCM rule is also consistent with the plasticity rule inferred from earlier physiological studies on familiarity effects in IT [9]. We developed a V1 circuit model with the BCM rule to successfully account for the familiarity suppression effects observed in the early visual cortex [8]. The model further predicts the sharpening of the tuning curves for the familiar stimuli in the early visual cortical neurons. Huang et al. [8] did not observe the tuning curve “sharpening.” However, in that study, only 25 familiar stimuli were trained and tested, which likely did not include the preferred stimulus of the neurons tested. In our simulation, we found that the tuning sharpening effect can be observed when 500 random natural images were used for familiarity training. Given that the familiarity effect is absent when the stimulus is limited to the receptive fields and the onset of the effect in V1 and V2 is 30 ms earlier than in IT [8], the development of sensitivity to global familiar images, 4-6 times larger than the size of the receptive fields of individual neurons, can be attributed to the formation of local recurrent circuits. These recurrent circuits exist in each visual area, processing familiar images in each area with progressively more global context down the ventral visual hierarchy.

The Hebbian-based cortical recurrent network in each visual area resembles a Hopfield network, with relatively local connections, capable of encoding associative memory. The connections in V1 and V2 extend outside the receptive fields but are as global as a Hopfield net. In addition to Hebbian excitatory horizontal connections, this cortical network is characterized by an additional strong surround iso-orientation (or iso-feature) inhibition and competitive local normalization, which is important for maintaining the stability of the network and for generating familiarity suppression and tuning curve sharpening effects. A neuron strongly activated by a specific familiar image, boosted by the excitatory horizontal circuit for encoding that image, would suppress other less-responsive neurons in the local neighborhood, leading to the familiarity suppression effect at the population level. Here, the super-linear activation function is used to model the neurons [25], but this specific mechanism is not necessary as familiarity effects can also be observed when a rectified linear activation function (ReLU) is used. Nonetheless, the supra-linear activation mechanism enhances the sparsity of the population responses as it more strongly suppresses low activity and amplifies high activity.

While recurrent circuits have been considered as a form of associative memory, we advanced the idea that the recurrent cortical circuit is performing a manifold transform, and training would induce modifications in this manifold transform. In general, a recurrent circuit is performing a globally nonlinear but locally linear manifold transform, in contrast to a single linear transform for all the points on the manifold used in traditional manifold learning [15, 16]. It is due to the nature of recurrent dynamics that in the vicinity of attractors, the behavior of the system is approximately linear, where the tuning derivatives can be untangled and decomposed into different components that fall along different dynamic modes operating on distinct timescales. In the locally linear manifold transforms, the slow mode of the network dynamic should be rotated during the training process to become more orthogonal to the dimensions of irrelevant variations of the stimulus concepts, so that their impact on the discriminative dimensions can be minimized.

We empirically connected familiarity training and manifold transform by showing that recurrent circuits trained with familiarity training perform the proposed manifold transform for different image concepts in the familiarity association experiment. We found that familiarity training decreases the distances in the noise direction relative to the distances in the stimulus direction locally around each fixed point, which is the expected behavior of manifold transform, resulting in a more robust representation of the global image concepts. We further confirmed that such behaviors were mediated by orthogonalizing the slow modes with the direction of irrelevant variation, as predicted by the theory. Thus, the nonlinear computation of the recurrent circuits shaped by familiarity training can be considered to be “thinking nonlinear globally and acting linearly locally” to achieve flexible manifold transformation around each remembered concept. These findings suggest that familiarity learning simultaneously sparsify the neuronal tuning (which leads to the suppression of population activities) and rotates the slow dynamical mode around each attractor (which results in the manifold transform). A natural question arises: why do these two phenomena co-occur in the learning process? One possibility is that, for the flexibility of each local manifold transform to rotate the slow modes against the specific irrelevant (*θ*) directions around each concept, the representations or memories should have less interference with each other. Thus, the sparsification of the representation is favored [40]. Nonetheless, future research should address this question by theoretically analyzing the causal relation between the sparsification and mode rotation during the learning medicated by the BCM rule.

This study has additional limitations that should be addressed in future research. Firstly, the architecture of our model does not follow the detailed biological neural circuit structure with different types of neurons and connectivity. Rather, we incorporate the standard circuit mechanisms of the macaque’s primary visual cortex, such as surround excitation/inhibition, normalization, etc., to construct a generalizable circuit model and assume such architecture design is canonical and applicable to the familiarity effects observed in V2, V4, and IT in principle. Future research should investigate how incorporating more biologically realistic neural circuit structures, including diverse neuron types and specific connectivity patterns, might influence the model’s explanatory power for the familiarity effects and capacity for the manifold transform. Another important remaining question is the broader applicability of the proposed manifold transform mechanism in different continuous transformations. While our current work demonstrates the effectiveness of this approach in handling noise corruption, its adaptation to other transformations, like rotation, translation, contrast modulation, and spatial frequency alterations, as well as to dynamic stimuli that vary in the temporal dimension, remains to be examined. These generalizations might require some variations at the circuit implementation level but should validate the generality of our proposed mechanism and provide insights into the fundamental principles of neural information processing of sensory stimuli.

## 4 Methods

### 4.1 Parameter setting and assumptions

The network we used is a standard V1 network with sparse feedforward encoding. The feedforward response is computed using *N*_*k*_ = 64 convolutional filter is obtained using the convolutional sparse coding algorithm that jointly minimizes the reconstruction loss and *L*1 norm of the per-hypercolumn activation [23]. The strategy improved the coding efficiency and was shown to enhance the characterization of the receptive field of neurons in the primary visual cortex [41]

Most of the parameters used in the model are rather standard. The time constant of excitatory neurons (*τ*_*e*_ = 20) double that of inhibitory neurons (*τ*_*i*_ = 10) [42]. The local extent of the surround inhibition and excitation is also relatively well-known based on the spatial extent of the surround inhibition [43], here we set the radius for mutual facilitation among excitatory neurons to be *R*_*e*_ = 2, and the radius for iso-orientation suppression is *R*_*i*_ = 1. Also, in Fig S1, we showed that the exact profile of the inhibition strength (uniform or Gaussian) is inconsequential in our model. The initial weight of plastic excitatory-excitatory connections is *w*_*ee*_ = 5. The absolute strength of the inhibitory surround *w*_*ie*_ is the main free parameter that was tuned for the model stability. We performed a parameter sweep (Fig S2) to understand the impact of this parameter on the main effects (sharpening of tuning curves and familiar suppression). In the main experiment, we typically set *w*_*ie*_ = 30. We set the synaptic time constant of BCM learning to be *τ*_*w*_ = 2*e*9, and the time constant of the exponential moving average for the BCM threshold to be 2*e*7.

### 4.2 Familiarity training for the familiarity effect

We trained the network on 500 natural images from CIFAR100. The size of the image is 32 ×32. Each image was convolved by 64 filters of size 9 ×9 derived from the convolutional sparse coding [23], resulting in outputs of size 8× 8, which is the feedforward input to excitatory neurons in the network. We set the network size *N*_*r*_ = *N*_*c*_ = 8 accordingly. Each image was shown to the network for 150 ms in one epoch of training. The network was trained for 5 epochs. The network was reset to the initial state (***r***^***e***^ = **0**; ***r***^***i***^ = **0**) between each image to mitigate inter-stimulus interference. During training, we scale the feedforward input by 30 to increase the training speed.

The initial BCM threshold for each neuron is determined by averaging its responses across all time steps and stimuli in the pre-trained network.

### 4.3 Quantification of familiarity effects (Fig 2, S1 Fig, S2 Fig, S3 Fig)

The familiarity suppression index is calculated as follows:

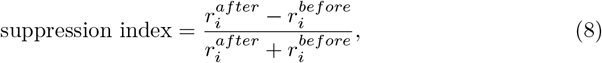

where 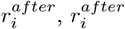 denote the steady-state response of neuron *i* after and before familiarity training. The neuron response was min-max normalized before computing the familiarity suppression index.

The sparsity index of a neuron tuning curve is introduced in [44], which is expressed as:

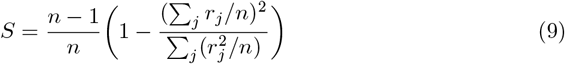

where *r*_*j*_ indicate the neuronal response to *j*^*th*^ stimulus. This index measures the tail length of the neuronal tuning curve; a high value indicates the tuning curve has a long tail, with only a few neurons having non-zero firing rates [45].

### 4.4 Derivation of locally linear manifold transform in the recurrent circuit (Fig 3)

Locally linear manifold transform is determined by dynamical modes derived from the Jacobian matrix of the linearized system. To obtain the Jacobian matrix and the dynamical modes, we first rewrite the network dynamic (Eq 1&2) into the matrix form:

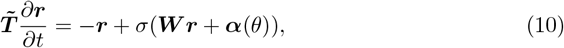

where 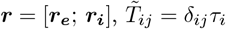,and *W* = [*W*_*ee*_, *W*_*ei*_; *W*_*ie*_, **0**] (We use “,” to separate column, “;” to separate row). ***α***(*θ*) denotes the sparse code input of *θ*. To perform linearization on Eq 10, we expand the network around the fixed point ***r***(*θ*) using Taylor series:

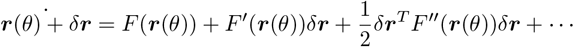

Since ***r***(*θ*) is the fixed points, *F* (***r***(*θ*)) = 0, and the second order term vanished assume small *δ****r***, we can obtain the linearized dynamic of the network around ***r***(*θ*) by substituting back Eq 10:

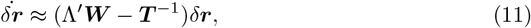

where 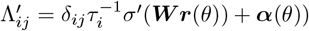.This matrix quantifies the sensitivity of each neuron’s firing rate to small changes in its membrane potential.

Eq 11 defines a simple linear system that approximates the dynamic of the original system near the fixed point ***r***(*θ*). We define Jacobian *J*(*θ*) = Λ^*′*^***W*** − ***T*** ^−1^. Note that the dependency of the Jacobian on the stimulus variable will be omitted in the notation for simplicity. Dynamical modes of the linear system can be found by diagonalizing the Jacobian using eigendecomposition. After diagonalizing *J*, the dynamic can be written as:

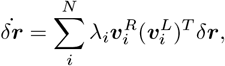

where 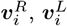 denotes *i*^*th*^ right/left eigenvector, and *λ*_*i*_ refers to the corresponding eigenvalue. Due to the orthogonality between left and right eigenvectors 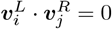,by projecting *δ****r*** onto the left eigenvector, we efficiently ‘diagonalize’ the dynamics, such that it is decoupled into multiple independent processes:

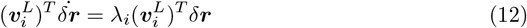

We define the dynamical mode as the left eigenvectors ***v***^*L*^, and the dynamical time scale as *τ* = − 1*/Re*(*λ*). Eq 12 shows that each dynamical mode describes a specific deviation pattern of population activity from the fixed point. The evolution of each pattern is independent and exponential, with the speed being controlled by the corresponding time scale *τ* . A negative *τ* entails escaping from the fixed point, while a positive *τ* entails decaying back to the fixed point.

The tuning derivative is computed by differentiating the steady-state response ***r***(*θ*) = *σ*(***W r***(*θ*) + ***α***(*θ*)) with respect to *θ*. After plugging in the eigendecomposition of the Jacobian, we obtain Eq 6

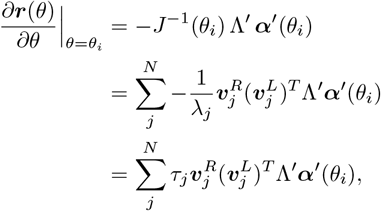

Again applying the orthogonality of the left and right eigenvectors, projecting the tuning derivative on the dynamical mode obtains:

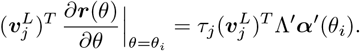

Projecting onto multiple dynamical modes results in Eq 7

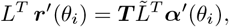

where the matrix L has columns 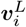, while 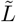 has columns 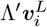.The diagonal matrix ***T*** contains the time scales *τ*_*i*_ of each mode on its diagonal.

### 4.5 Familiarity training for the familiarity association experiment

For the familiarity association experiment, we randomly selected five 32×32 images from the CIFAR-100 dataset to serve as the target images. The images were converted to the grey scale before adding the noise. To corrupt the target image *l* by noise level *n*%, we randomly chose *n*% pixels and sampled the pixel value from *ε* ∼ 𝒰 (0, 1) to replace the original one. This results in a distribution of stimuli ***x***_*n*,*l*_ ∼*p*(***x*** | *n, l*). We sampled 3 noise levels (10%, 30%, 50%) and 10 stimuli for each noise level and target image; hence, there are 5 × 3 × 10 target-noise pairs in total. The feedforward input was obtained by convolution with the same sparse coding filter bank as in 4.2.

The network architecture was configured with dimensions *N*_*r*_ = *N*_*c*_ = 8 to match the image size. The network was trained for 5 epoches. During each training epoch, the noisy image was first presented to the network for 150 ms, then the clear image for 150 ms. To ensure the specificity of training on the presented image, the network state was reset to ***r***^***e***^ = **0**; ***r***^***i***^ = **0** prior to the presentation of each image. During training, we multiply the feedforward input by 30 to accelerate the training process. The initial BCM threshold for each neuron was established by calculating the mean of its responses across all time steps and stimuli in the pre-trained network.

### 4.6 Fixed points analysis in the familiarity association experiment (Fig 4, S3 Fig)

The fixed point was found by forward simulating the network until convergence in the probe test period. The trajectory was visualized by reducing the dimension to 3 via PCA. The fixed point manifold was visualized by embedding the manifold in 2D via Multidimensional Scaling (MDS) that preserves the pairwise Euclidean distances between fixed points of interest.

The relative distance is computed as:

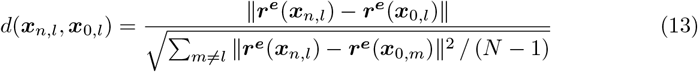

where *N* is the number of target images, and ***r***^***e***^(***x***_*n*,*l*_) refers to representation of ***x***_*n*,*l*_ which was sampled from the conditional distribution *p*(***x*** | *n, l*) (where *n* denotes the noise level and *l* denotes the target image). We sampled 10 stimuli for each *n* and *l*, and the relative distance was averaged over those 10 stimuli.

To measure the directional alignment, we computed the cosine similarity given by

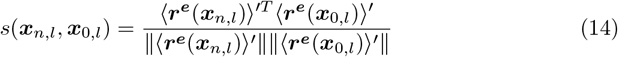

where ⟨***r***^***e***^(***x***_*n*,*l*_) ⟩ refers to the averaged representation of 10 stimuli sampled from *p*(***x*** | *n, l*)⟩. ⟨***r***^*e*^(***x***_*n*,*l*_) ^*′*^ = ⟨***r***^***e***^(***x***_*n*,*l*_)⟩ −⟨***r***^***e***^(***x***_0,*l*_)⟩ _*l*_, which denotes the population vector of excitatory neurons that starts from the mean of target images.

### 4.7 Analysis of the dynamical modes in the familiarity-trained network (Fig 5, S4 Fig)

To analyze dynamical modes and their associated decaying time constants in our neural circuit model, we forward-simulated the system until it converges to find the fixed point and used the numerical method to obtain the eigenvalues and eigenvectors of the Jacobian matrix. We define *L* to be the matrix of eigenmode, whose columns are top M slow left eigenvectors 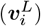 of the *J*(*θ*) (M ranges from 1 to 200). Similarly, 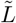 is column concatenation of 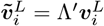.The decaying time constant of mode *i* is: *τ*_*i*_ = −1*/Re*(*λ*_*i*_), where *λ*_*i*_ is the *i*^*th*^ eigenvalue of *J*. Eigenvectors and eigenvalues were pooled across visual concepts, noise levels, and training epochs. The mean and s.e.m. of *τ*_*i*_ and the mean of 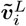 were plotted.

Due to the complexity and unobservability of the feature space of the natural images, we used the distance between the nearby fixed points to approximate the derivative.

Specifically, when the linearization was performed at the representation of stimulus ***x***_*n*,*l*_ sampled from the conditional distribution *p*(***x*** | *n, l*) (where *n* denotes the noise level and *l* denotes the visual concept), the noise distance is:

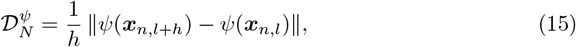

where *h* denotes the increment of noise level to the adjacent noise level, with a unit of 10%. In total 10 noise pairs (***x***_*n*,*l*_, ***x***_*n*,*l*+*h*_) were sampled from each conditional distribution (the linearization was always performed at ***x***_*n*,*l*_), and the distance was averaged over noise pairs. The concept distance is

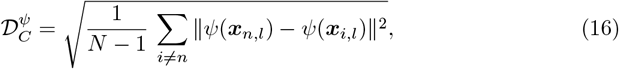

where *N* is the number of visual concepts. Similarly, ten concept sets (***x***_*n*,*l*_, *n* = 1, …, *N*) were sampled from each conditional distribution (the linearization was always performed at ***x***_*n*,*l*_), and the distance was averaged over concept sets. The normalized distance is computed as the ratio of noise and concept distance 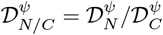,and the ratio was computed before averaging over noise pairs and concept sets. *ψ* can be input ***α***, representation ***r***^***e***^, input projection 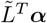 and output projection *L*^*T*^ ***r***, where *L* and 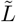 are derived from the Jacobian of ***x***_*n*,*l*_.

In Fig S4D-E, the relative difference between 𝒟_*N/C*_ in *ψ*_1_ and *ψ*_2_ were computed using the formula: 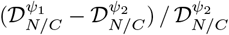 . To compute the correlation in Fig S4F-G, 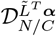 and 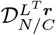 were concatenated across 3 epochs (pre-training, epoch1, epoch5), and the Pearson correlation coefficient is then computed between the concatenated distances and the corresponding epochs.

## Supporting information

**S1 Text Supplementray figures S1-S4**. Additional results of numerical experiments

## Acknowledgments

This work was supported by NSF CISE RI 1816568 and NIH R01 EY030226-01A1. We thank Wenhao Zhang and Carl Olson for their insightful discussions.

## Data availability statement

All code written in support of this publication is publicly available at https://github.com/leelabcnbc/familiarity-circuit-V1.git. Simulation input files and generated data are available from https://doi.org/10.5281/zenodo.14729086

