## Supplemental Figures for "Manifold Transform by Recurrent Cortical Circuit Enhances Robust Encoding of Familiar Stimuli"

### Supplementary Figures

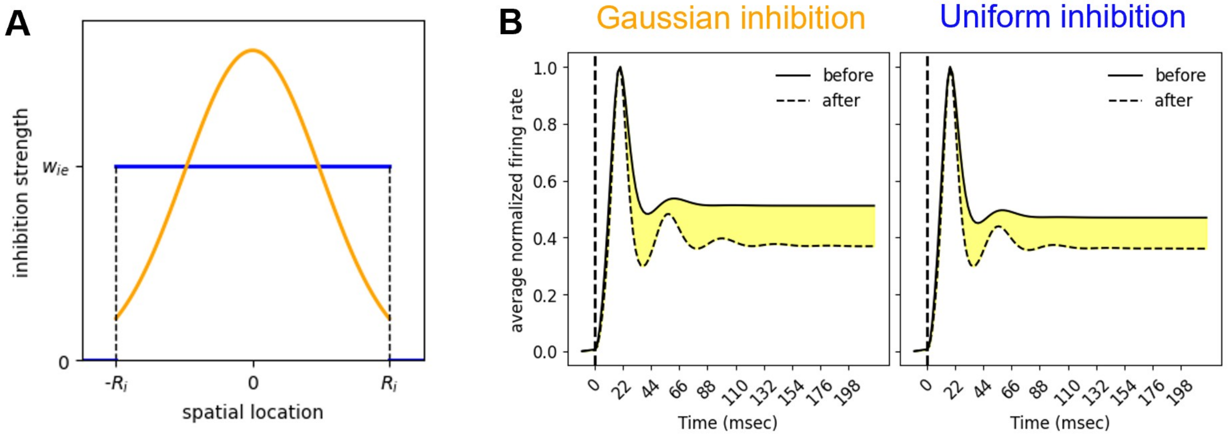


Figure S1: **Comparison between Gaussian inhibition and uniform inhibition.** (**A**) Illustration of Gaussian and uniform inhibition. The X-axis is the spatial location, where the target inhibitory neuron is at 0, and its inhibitory neighborhood spans the range from $-R_i$ to $R_i$. The Y-axis is the strength of $w_{ie}$. Blue line: uniform inhibition, where all E to I strength equals the initial weight $w_{ie}$. Orange line: Gaussian inhibition, where the E to I strength varies around $w_{ie}$ in a distance-dependent manner. (**B**) Familiarity suppression in models with Gaussian inhibition and uniform inhibition. The yellow area indicates the reduction after training. The familiarity suppression effect is not affected by the inhibition profile.

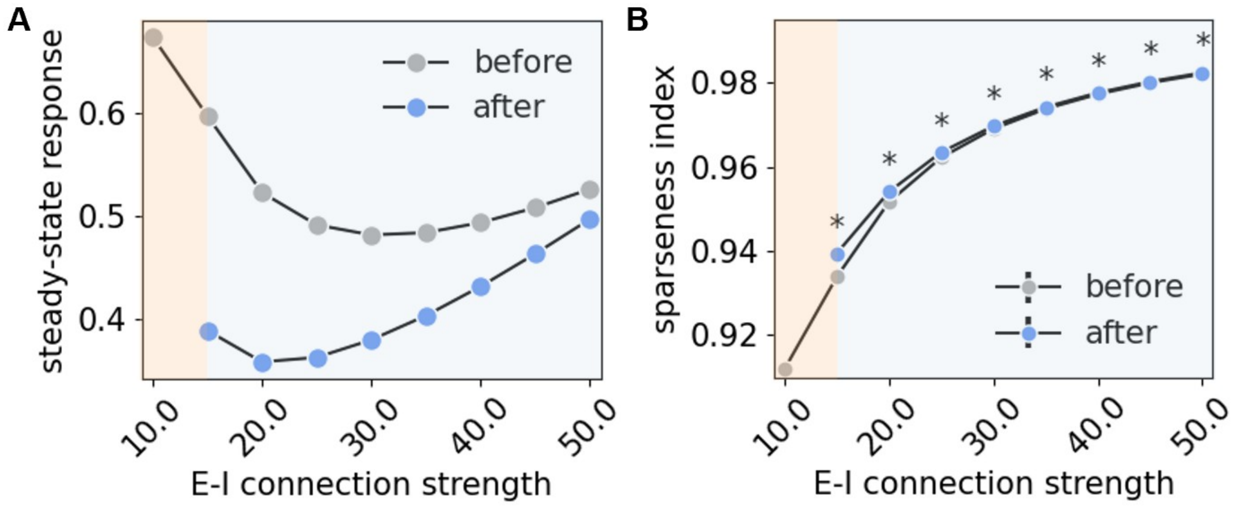


Figure S2: **Parameter sweep of $w_{ie}$.** (**A**) Effect of varying $w_{ie}$ on familiar suppression. Grey dots: before training; blue dots: after training. The suppression effect peaks at the critical points where the network transitions from the unstable to the stable regime, and the suppression effect decreases as $w_{ie}$ moves away from the critical point. (**B**) Effect of varying $w_{ie}$ on tuning curve sharpening. Grey dots: mean value before training; blue dots: mean value after training. * means after-training sparseness is significantly larger than before-training sparseness. Like suppression effects, the tuning sparsification also reaches the maximum at the critical points and decreases as $w_{ie}$ moves away from the critical point. In both **A** and **B**, the yellow area indicates the unstable regime and the bluw area indicates the stable regime.

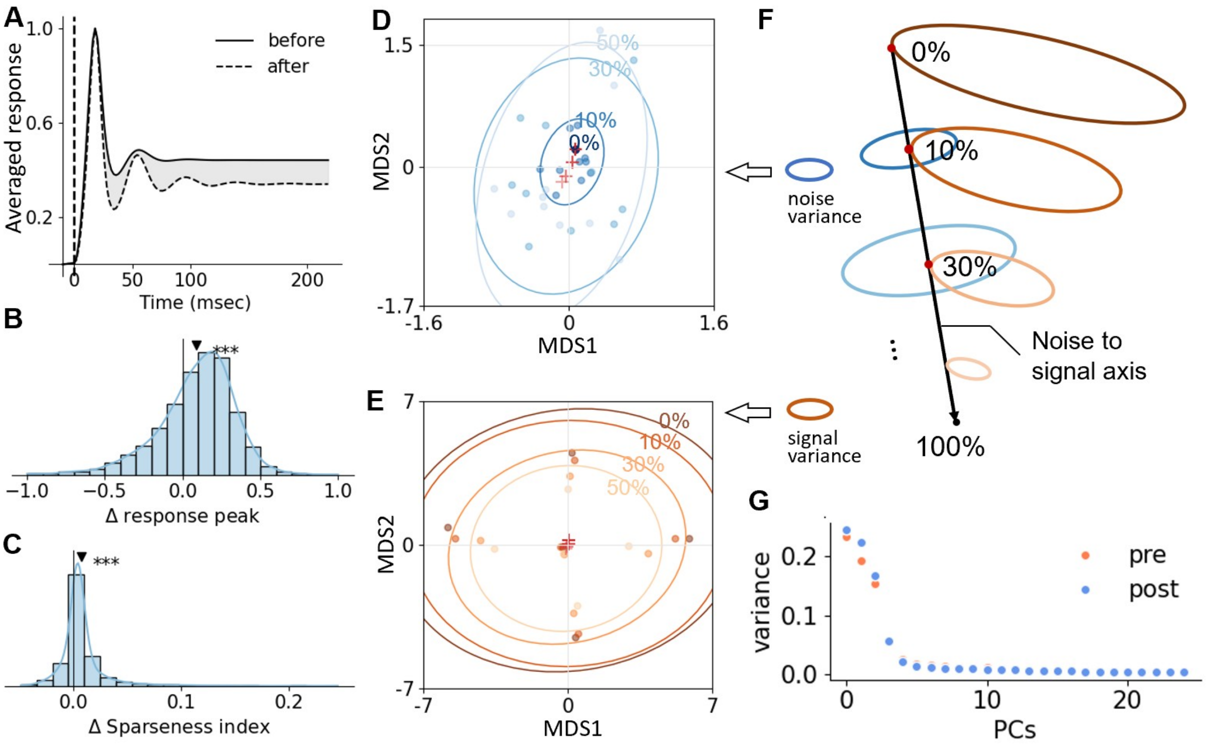


Figure S3: **Neuronal tuning and population activity in familiarity association experiment.** (**A**) Familiarity suppression effects were also observed in the association experiment. The grey area represents the reduction in the average steady-state response caused by familiarity training. (**B-C**) Tuning curve sharpening was also observed in the association experiment. The triangle marker indicates the mean of the distribution. ***: $p < 0.05$ means the mean of the relative difference ((after-before) / (after+before)) is greater than 0. (**D**) Visualization of the clusters of different noise levels averaged across different target images in epoch5 trained network. The figure represents a top-down view of the blue cone in **F**. Dots: noisy image samples. Red crosses: cluster means of averaged clusters. Ellipses: covariance of averaged clusters. Deeper color means a lower noise level. (**E**) Visualization of the manifold formed by all cluster means in epoch5 trained network. The figure represents a top-down view of the orange cone in **F**. Dots: cluster means. Red crosses: average of cluster means of the same noise level. Ellipses: covariance of cluster means of the same noise level. Deeper color means a lower noise level. (**F**) Illustration of the representation manifold. The blue and orange cones are visualized in **D** and **E**, respectively. (**G**) Explained variance of the first 25 principal components of the representation in the pre-training network (pre) and epoch 5 trained network (post). The dimensionality of the representation decreased after training.

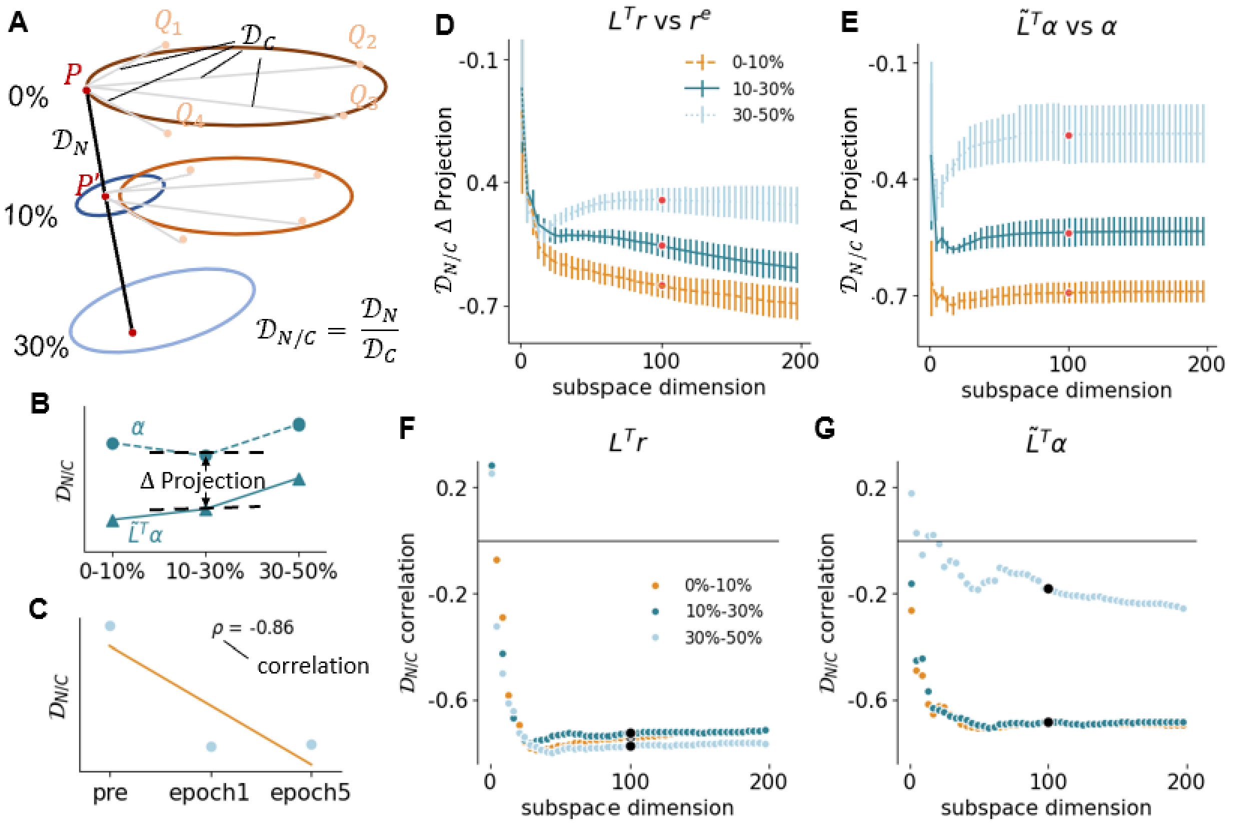


Figure S4: **Relative differences in distances and training effects with different projection subspace dimensions** (**A**) Illustration of normalized noise distance ($\mathcal{D}_{N/C}$), noise distance ($\mathcal{D}_N$), and concept distance ($\mathcal{D}_C$). When the network is linearized at fixed point $P$, $\mathcal{D}_N$ (0% to 10%) is proportional to the Euclidean distance from $P$ to $P'$, and $\mathcal{D}_C$ (0% to 10%) is the averaged Euclidean distance from $P$ to $M_1$, $M_2$, $M_3$, and $M_4$. $\mathcal{D}_{N/C}$ is the ratio of $\mathcal{D}_N$ and $\mathcal{D}_C$. (**B**) $\Delta$ projection in **D-E** is the percentage decrease in the normalized noise distance after projections. (**C**) Correlation in **F-G** refers to the Pearson correlation coefficient between normalized noise distance and training epochs (pre-training, epoch1, epoch5). (**D-E**) Percentage decrease in the normalized noise distance after projection (left: output projection; right: input projection) as a function of the projection subspace dimension. The effect of projection on the normalized noise distance is consistent across subspace dimensions. All distances are computed in the epoch5-trained network. The triangle marker and the black dots denote the distance values in the 100-dimensional projection subspace, which are results shown in figure 5. (**F-G**) Pearson correlation between training epochs and projected normalized noise distance (left: output projection; right: input projection) as a function of the projection subspace dimension. The training effects are presented in a wide range of subspace dimensions. The black dots denote the distance values in the 100-dimensional projection subspace, which are results shown in figure 5.
